# ZebraTrack: An Open-Source Object Detection Algorithm to Detect and Track Larval Zebrafish Motor Touch Responses

**DOI:** 10.1101/2025.10.17.683088

**Authors:** Adrien Lacroix, Gary A.B. Armstrong

**Affiliations:** Department of Neurology and Neurosurgery, McGill University, Montreal, QC, Canada

## Abstract

**Motivation:** Zebrafish (*Danio rerio*) are a model organism used for the study of vertebrate development, disease and drug discovery. Two-day old larval zebrafish exhibit burst swimming behaviour that can be elicited by a light touch to the tail. Larval motor touch-responses are frequently video recorded and later analyzed. Methods to robustly analyze these videos in a reproducible and time-efficient manner are reliant on manual tracking, which is prone to experimenter bias and error.

**Results:** Here we present ZebraTrack, a machine learning-based program, which employs Ultralytics’ YOLOv8 nano (YOLOv8n) object detection algorithm to automatically analyze larval touch response videos. The program breaks down video files into their constituent frames and passes these through a custom-trained YOLOv8n algorithm to detect the presence of a single larval zebrafish. ZebraTrack then refines the tracking data output by the model and tabulates it into an excel spreadsheet. The program then computes and extracts four relevant swim metrics: swim duration (s), swim distance (mm), mean swim velocity (mm/s), and max swim velocity (mm/s). ZebraTrack rapidly accelerates the analysis process, while also eliminating the errors associated with manual tracking. Furthermore, it allows for high-throughput analysis of larval touch response videos and can detect subtle differences in motor metrics arising as a result of temperature differences, demonstrating that utility of this tracking algorithm.

**Availability and implementation:** ZebraTrack is available for download at https://github.com/Armstrong-Lab-70/ZebraTrack.

## Introduction

Zebrafish are a widely used vertebrate model organism due to their rapid development, semitransparent embryo, and accessible nervous and motor systems for investigating behaviour (Stewart et al., 2014). Furthermore, the zebrafish genome shares a high degree of genetic conservation with the human genome and is amenable to editing including by use of the CRISPR-Cas9 system to generate both knockout and knock-in models (Varshney et al., 2015; Albadri et al., 2017; Simone et al., 2018), making them an excellent model for the study of biology. Moreover, their suitability for large-scale genetic and small molecule screens allows for the identification and functional analysis of genes and cellular pathways involved with disease. Among vertebrate models, zebrafish have emerged as a particularly valuable organism for investigating development and function of the motor system (Saint-Amant and Drapeau, 1998, 2000; Brustein et al., 2003; Masino and Fetcho, 2005), where motor touch responses can be used to evaluate the therapeutic potential of molecules (Patten et al., 2017; Goldshtein et al., 2020).

The zebrafish embryo develops the ability to respond to tactile stimuli at 21 hours post-fertilization (hpf) (Saint-Amant and Drapeau, 1998), where its emerging motor system changes from spontaneous coiling behaviour (Saint-Amant and Drapeau, 1998) to a propulsive behaviour required for hatching at two days of age (Kimmel et al., 1995). The touch-evoked escape response is frequently studied in two-day old larvae, where a light touch to the tail or head elicits burst swim behaviour that last a few seconds (Kimmel et al., 1995). The motor touch-response assay is conducted by placing a single larva into the centre of a large petri dish filled with system water, where burst swimming behaviour is evoked using a pair of forceps. Touch-evoked responses are recorded from above using a video camera, using both high resolution and frame rates enabling data to be of sufficient quality for *post hoc* manual analysis (Sztal et al., 2016). Video capture and analysis softwares such as StreamPix5 can be used for analysis of motor touch-response assays in order to accelerate the analysis process (Sztal et al., 2016). However, high quality cameras and analysis softwares designed for laboratory use can be expensive and inaccessible to many researchers. Free alternative softwares, such as Softonic for video capture and ImageJ for analysis, provide an accessible yet imperfect solution (Sztal et al., 2016). Manual tracking using ImageJ requires frame-by-frame analysis and data post-processing (*i.e.*, extraction of relevant swim metrics from tracking data) to be performed by hand, where it is time consuming and prone to errors.

In this study we developed a program named ZebraTrack, with the objective of creating a tool for rapidly and reproducibly analyzing larval zebrafish motor touch-response videos. This program was developed using the Python programming language on Google’s Colaboratory platform, making the code and ZebraTrack program freely accessible to investigators. This program is centered around Ultralytics’ YOLOv8n object detection model, which is an open-source computer vision algorithm available as a Python package (Varghese and Sambath, 2024). Recent advances in object detection algorithms provide a rapid, accurate and reliable analysis tool for tracking in videos (Sohan et al., 2024). In our application, the YOLOv8n algorithm is given individual videoframes of two-day old larval motor touch-response videos in order to predict the location of a single larval zebrafish in a sequence of video frames. Then, it refines the tracking data using a custom computer vision algorithm before calculating swim metrics. ZebraTrack generates an excel spreadsheet containing four swim metrics: swim duration (s), swim distance (mm), mean swim velocity (mm/s) and maximum swim velocity (mm/s). ZebraTrack is capable of rapidly and unbiasedly analyzing multiple videos, enabling researchers to carry out accurate high-throughput data analysis of larval motor patterns.

## Materials and Methods

### Zebrafish husbandry

Adult zebrafish (*Danio rerio*) were maintained and bred according to standard protocols under a 14/10 light/dark cycle at 28.5 °C (Westerfield, 2000). All experiments were performed in compliance with the guidelines of the Canadian Council for Animal Care and performed at the Montreal Neurological Institute at McGill University (Montreal, Quebec, Canada) and approved by the Animal Care Committee (AUP#: MNI-7890).

### Larval touch-responses

The touch-evoked motor response assay was performed as previously described (Armstrong and Drapeau, 2013). Larvae were placed in the center of a circular dish (150 mm diameter) containing system water at approximately 1 cm in depth. Burst swimming behaviour was evoked by a brief touch to the tail and the swim pattern was recorded from above at 30 Hz using a digital camera (Grasshopper 2 camera; Point Gray Research). Swim duration, distance, mean and maximum velocities were quantified using the manual tracking plugin for ImageJ. For the purposes of quantifying YOLOv8n model tracking performance, these metrics were also analyzed based on tracking data obtained automatically through the model’s predictions.

### Statical analysis

All statistical comparisons were performed using Prism 10 (GraphPad Software Inc.). Data is presented as mean ± standard error. A Shapiro-Wilk test was used to assess normality. When data were normally distributed, a one-way ANOVA followed by Tukey’s multiple comparisons test, or Welch’s t-test were performed to identify were differences lay. When data were not normally distributed, a Kruskal-Wallis test followed by Dunn’s multiple comparisons test, or Mann-Whitney U-test were performed to identify were differences lay. Significance was assessed at *p* < 0.05.

## Results

### Workflow for motor touch response tracking and analysis

The automated tracking program is designed to run on Google’s Colaboratory platform (URL: https://colab.research.google.com/) and requires that users have a Google Drive account in order to upload videos. The program analyses grayscale motor touch-response videos of individual 2 day-post fertilization (dpf) larvae filmed at a frame rate of 30 Hz (or greater) and with a resolution of 480 by 640 pixels. The program begins by prompting users for a frame rate and an x-y calibration (x-y cal) value to relate pixel space to real space, and functions under the assumption that these two values (frame rate and x-y cal) remain constant throughout all videos in the video folder specified by the program. The program checks for erroneous user input before commencing video analysis using a custom-trained version of Ultralytics’ YOLOv8n object detection algorithm. Then, ZebraTrack analyzes each video, where an output comma-separated values (.csv) file containing tracking predictions is generated for each video. As the algorithm can make erroneous predictions, ZebraTrack also attempts to rectify any mistakes that may have been generated by the prediction process. To further accelerate analysis, the program also creates an excel (.xlsx) file. This file contains two sheets; one containing the tracking data from each video in the folder, and another which is a summary sheet presenting four swim metrics for each video. Finally, the program also provides tracking data in individual .csv files compatible with ImageJ’s manual tracking plugin for manual review.

### Data collection and extraction

Larval touch-response tracking data (n = 1084 videos) collected from previous projects were repurposed for the creation of a training dataset. These data included .xlsx files containing tracking data, as well as corresponding videos from which the tracking data were manually obtained in previous research. Custom python scripts were written to extract data from .xlsx files into a YOLOv8-compatible annotation format (**Fig. 2A**). Another custom python script was written for the purpose of extracting individual frames of video and matching them to the corresponding annotations (**Fig. 2B**).

**Figure 1.**
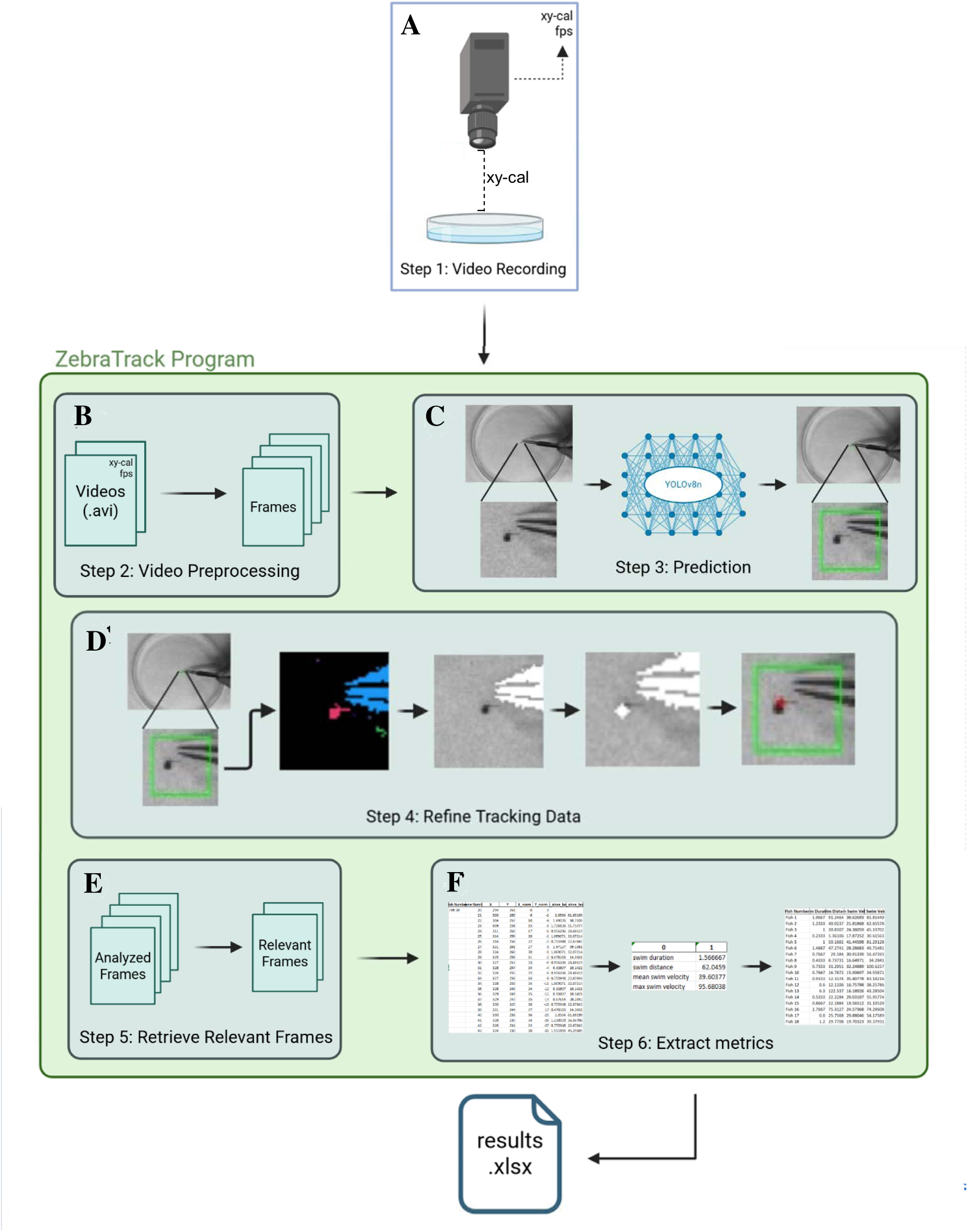
ZebraTrack application workflow. **A**, Users must are prompted to input a folder which contains videos of touch-evoked motor responses of individual larvae. Users are required to provide an x-y calibration value, obtained by dividing the width of the camera frame in meters by the width of the camera frame in pixels. This calibration value allows the tool to relate pixel distances to real-life distances and must be adjusted every time the camera’s height is adjusted. Users are also required to provide the frame rate at which the videos were recorded. The program assumes that both the x-y calibration and the frame rate values remain the same throughout a batch of input videos. **B**, ZebraTrack splits recorded video sequences into individual frames for the YOLOv8n model. **C**, Predictions are obtained from the model. The raw tracking data is output in the form of a bounding box, which identifies an area within which the model predicts the larva to be present. **D**, ZebraTrack applies computer vision techniques (through the OpenCV Python library) to refine the tracking data. When necessary, ZebraTrack identifies and removes distractor objects. First, the individual frame is segmented into a series of connected components. Second, the largest connected component is identified, then masked and removed from the image. This operation is performed to remove the forceps from the image. Third, the program takes the average location of the darkest pixel across all frames of a video, then masks and removes that point. This operation is performed to remove the dark, central dot marked on the petri dish. Fourth, and finally, the location of the larval head is identified as the darkest point in the bounding box. This process is repeated individually for each frame of video. **E**, The refined tracking data is processed to detect the start and end of the burst swimming behaviour. **F**, After the tracking data has been refined and narrowed down to relevant frames, it is processed into 4 metrics; swim distance, swim duration, mean swim velocity and maximum swim velocity into an excel file containing tracking data as well as swim metrics for each video.

**Figure 2.**
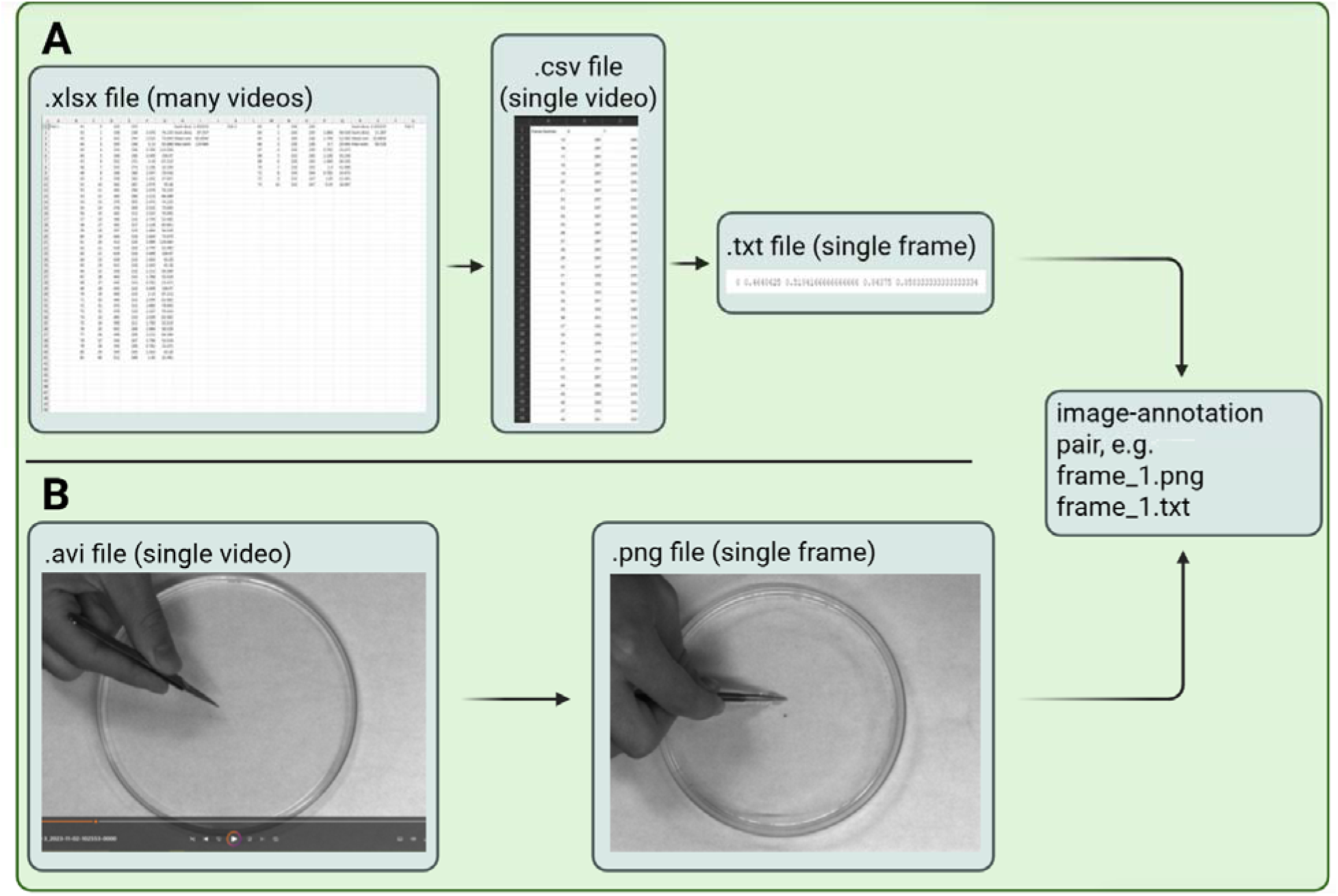
Data processing pipeline for dataset curation. **A**, The process begins with matching excel files to corresponding videos. After manual renaming of files, a custom python script extracts image-annotation pairs from the manual tracking data found in the excel files and their corresponding larval touch response videos. Each excel file is broken down into individual .csv files, each representing a single video. Then, the .csv files are broken down into individual text files in the YOLOv8 PyTorch .txt format. **B**, Following this, the video files (.avi file format) are broken down into individual images, each named correspondingly to their matching annotation file allowing for the training using YOLOv8n.

### Training YOLOv8n

The final dataset used for training comprised 65,416 image-annotation pairs with annotations provided in the YOLOv8 PyTorch .txt format. A train-test-val split (80-10-10) was applied to separate the dataset into training, testing and validation sets. Only the training and validation sets were provided to the model during training. The pretrained YOLOv8n model was selected based on shorter training time and the relative simplicity of the detection task, since larger pretrained models would require additional computational power from the GPU to train for a minimal increase in training performance. To train the model we used a NVIDIA GeForce RTX 2070 Super (8192mb) GPU, for 400 epochs, and a patience of 10 epochs to avoid overtraining. An epoch is defined as a complete pass through the entire training dataset within the context of a larger training session. Therefore, by setting the training to take place for 400 epochs, the model will complete 400 passes through the entire training dataset. In addition, patience is a hyperparameter which defines the number of epochs for which training metrics must stagnate before training is halted prematurely. This prevents overfitting, which occurs when the model learns to perform well on the training dataset at the expense of generalizability to new data samples. To optimize training speed, batch size and number of workers were set to 8 and 4 respectively. All other model hyperparameters were left at default values.

### YOLOv8n algorithm

The model used to develop our larval tracking program was Ultralytics’ YOLOv8n (Varghese and Sambath, 2024). While the performance of the nano model is limited due to its relatively low number of parameters (2 million), it is still well-suited for simple object detection tasks which must run in computationally constrained environments. As such, this model provides an accessible object detection algorithm which performs well without requiring expensive computer hardware. Briefly, the algorithm can be broken down into three modules: a backbone, a neck, and a head (Sohan et al., 2024; Varghese and Sambath, 2024; Yaseen, 2024). The backbone module is constituted of a convolutional neural network which extracts multi-scale features from input images (Sohan et al., 2024). The neck module is a multi-scale feature integrator, which refines the features extracted by the backbone and fuses them in a process which is required for detecting objects of varying shapes and sizes (Sohan et al., 2024). The head module generates the final predictions, including class labels, confidence scores and bounding box coordinates for each detected object (Sohan et al., 2024).

### YOLOv8n performance with varying training dataset sizes

Although there are many measures of performance in object detection models, mean average precision (mAP) is frequently used as a metric to describe overall performance (Chumachenko et al., 2022; Zaidi et al., 2022; Talha et al., 2023). This metric is preferred because it encapsulates many other metrics, such as precision, recall and intersection-over-union into a single value (Chumachenko et al., 2022). Precision is measured as the number of true positives over the total number of true positives and false negatives, which measures what proportion of the predictions are correct (Chumachenko et al., 2022). Recall is measured as the number of true positives over the total number of true positives and false negatives, which measures what proportion of the total possible correct predictions were obtained (Chumachenko et al., 2022). Intersection-over-union (IoU) measures, for each prediction, the overlap of predicted and ground truth bounding boxes (Chumachenko et al., 2022). Ultralytics’ mAP50-95 metric encapsulates both precision and recall, at an intersection-over-union (IoU) threshold of 50 to 95 percent, across all classes (Varghese and Sambath, 2024). However, since there is only one class of interest (zebrafish larva) for our model, the mAP metric can be viewed as equivalent to average precision (AP) alone. The mAP metric is calculated by finding the area under the precision-recall curve, and mAP50-95 refines this by calculating the average mAP at 10 separate IoU values ranging from 0.50 to 0.95, thus giving a comprehensive overview of the model’s performance in a single metric (Varghese and Sambath, 2024).

As the data were collected over the course of the project, the model was trained several times using datasets of increasing size. The performance of the model using the first dataset of 18,189 images achieved a mAP50-95 value of 0.74. When the size of the dataset was increased to 20,014 images, the model attained a mAP50-95 value of 0.79. However, on the final dataset of 65,416 images, we observed a decrease in performance to an mAP50-95 value of 0.74. These mAP50-95 values were obtained by utilizing Ultralytics’ built-in “.val()” method, specifying the custom-trained model version as well as the dedicated test dataset, which was novel to the model. By using the test dataset, which constitutes 10% of the entire dataset, we ensured that the obtained performance metrics are representative of the model’s performance on unseen data, rather than data used during training such as the training and validation datasets.

### Prediction generation and refinement

After receiving the appropriate user input, ZebraTrack reads through each video frame-by-frame, feeding each frame to the custom-trained YOLOv8n algorithm. When the algorithm generates a prediction, it outputs coordinates for a bounding box, which delineates a square region in which the model predicts the larva will be present. The algorithm outputs predictions for each frame, which are then saved in a .csv file that represents all the predictions for each frame of a given video. Once all videos have been analyzed using YOLOv8n, the program refines the predictions. As YOLOv8n outputs bounding boxes that may center on random points along the larvae’s body, ZebraTrack was designed to use a simple computer vision (CV) algorithm to refine the models’ predictions. This simple CV algorithm works to: (1) detect and mask a dark central dot if present within the bounding box region; (2) detect and mask forceps used to evoke motor touch-responses, if present in the bounding box region; and (3) find the local minimum pixel brightness value in the bounding box region (which is assumed to be the larva’s head). This simple algorithm works to refine the bounding box predicted by YOLOv8n to a singular cartesian point within the image which represents the location of the larva’s head. Briefly, the logic of the algorithm is based on heuristic assumptions, which rely on the highly standardized nature of touch response videos. It assumes that the larva’s head is the darkest point in each bounding box, which is true except for in two cases. The first case involves the presence of a dark, central dot on the petri dish that researchers use to initially position larvae in the centre of the dish. To determine the location of the centering dot, the algorithm performs a search for the darkest point in every video frame and applies a mask to the average location of the darkest pixel. Importantly, the program functions under the assumption that there will always be a central dot, as is the case for all the videos the YOLOv8n algorithm was trained on. The second case involves the presence of forceps, used to evoke motor behaviour by lightly touching the larval tail. The algorithm applies a function from the OpenCV python library to detect the largest connected component currently in frame and applies a mask to this object. In cases where both the dot and forceps are present, the algorithm applies both masks. In cases where the position of the larva’s head cannot be reliably determined, the default is value is the center of the bounding box. Later steps in the program review the predictions in order to remove erroneous predictions.

### Prediction correction

While the custom-trained YOLOv8n model performs well on larval zebrafish detection tasks, it can make prediction errors. For example, the model may detect two larvae when only one is present in a frame (note: ZebraTrack can only be used to track single larva touch-responses), or it may erroneously predict the single larva’s position, or fail to generate any predictions for a given frame. While these errors are uncommon, ZebraTrack addresses them by searching through each .csv file generated by the prediction step. This serves to detect errors through three main approaches. (1) If the program finds failed predictions in the first few or last few frames, it removes them from the .csv file, as these frames usually do not contain larval movement. (2) It verifies that each velocity value is realistic; if a larva’s velocity has exceeded 300 mm/s over the last frame, the prediction is deleted and replaced with an empty value. (3) If empty values are found, after the previous two steps have been completed, they are assumed to be failed detections and are linearly interpolated. This is achieved by linearly interpolating the larva’s position using the previous frame and following frame, as this preserves a sufficient amount of information about the larva’s swim pattern. Since the error correction process functions under a number of assumptions, it is possible that ZebraTrack can fail to properly address errors. ZebraTrack thus also generates additional .csv files which contain the predicted locations of the larva in each frame for each video. These .csv files have been formatted to be compatible with ImageJ’s “manual tracking” plugin, allowing users to manually review ZebraTrack’s output for errors.

### Extraction of relevant frames

Given that the YOLOv8n algorithm functions solely to locate objects within an image, the machine learning algorithm itself cannot determine when a fish’s movement pattern begins or ends. As such, ZebraTrack takes a simple thresholding approach to detect the start and stop of larval swim patterns. To detect the start of a burst swim pattern, ZebraTrack applies a One Euro filter to the predicted positional data and marks a threshold for movement initiation at 30 mm/s. The One Euro filter is designed for smoothing time series data. It functions as an adaptive low-pass filter, adjusting its smoothing strength based on the rate of change in the signal (*e.g*., position of the larval head) (Casiez et al., 2012). When the signal changes slowly, the filter applies stronger smoothing to reduce noise; whereas when the signal changes rapidly, it decreases smoothing to preserve responsiveness. To achieve this dynamic filtering, the One Euro Filter applies a standard exponential moving average with a dynamic cutoff frequency that varies according to the signal’s derivative (Casiez et al., 2012). To detect the end of a swim pattern, we apply the One Euro Filter to the positional data and calculate a velocity value. When the larva’s velocity dips below the threshold value of 15 mm/s, the swim pattern is considered terminated. While the thresholding approach described here functions robustly, it can malfunction and result in erroneous start and stop times. For example, we conducted an analysis on data obtained manually versus data obtained by ZebraTrack and observed a slight trend of prematurely detecting the beginning of swim patterns (**Fig 3A**). In contrast, the program performs robustly at detecting the end of swim patterns (**Fig. 3B**).

**Figure 3.**
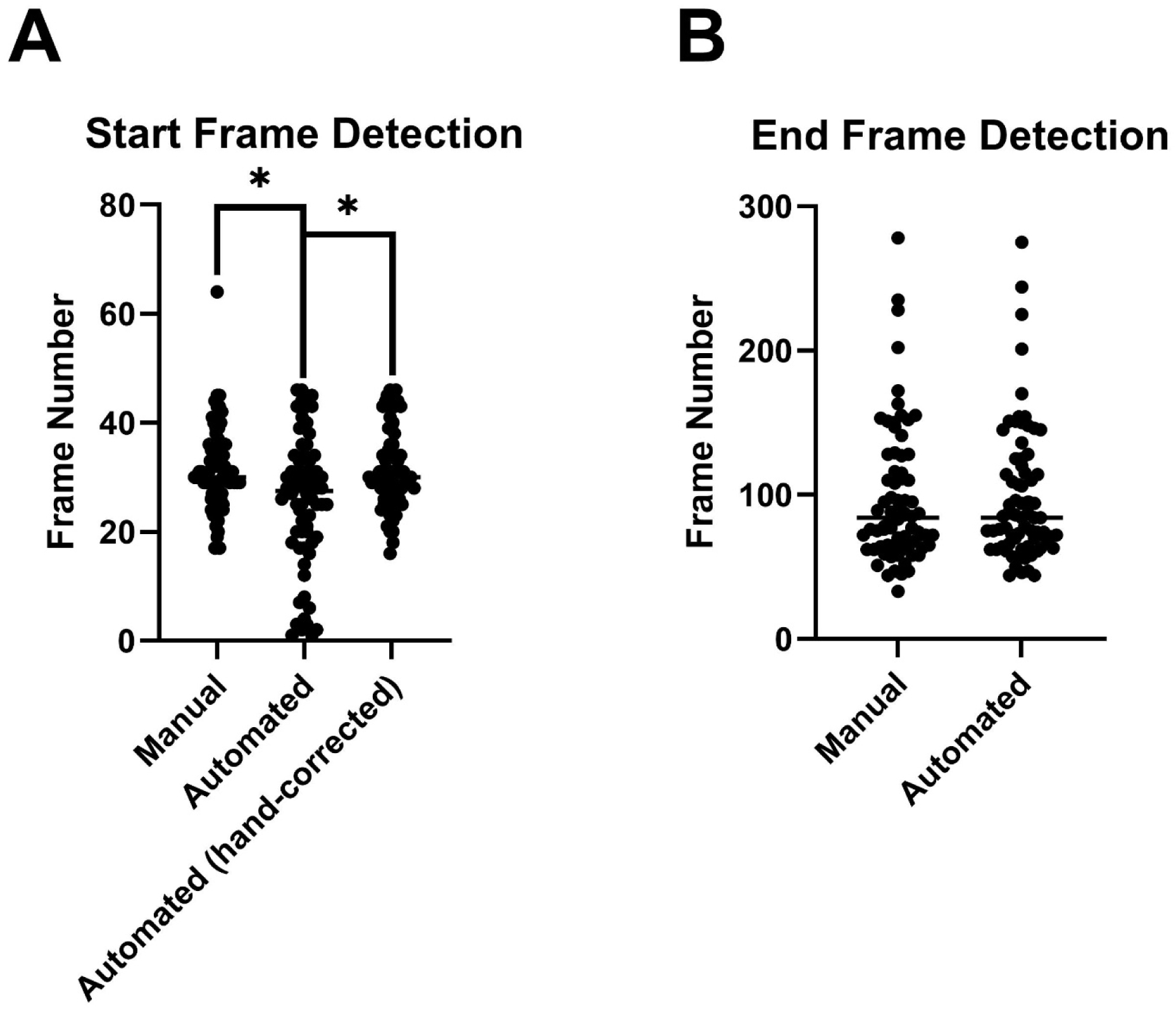
ZebraTrack adequately detects the start and end of larval zebrafish swim patterns. **A**, ZebraTrack program tends to prematurely detect the start of larval swim patterns. The manual group shows the frames at which the start of a swim patterns were detected, with each datapoint representing one video. Automated detection is significantly different from manual detection of swim pattern starts (*p* = 0.016), but with manual correction the data is not significantly different. However, manual correction of the data output by ZebraTrack can be performed to rectify these errors. **B**, ZebraTrack program accurately detects the end of larval swim patterns, as evidenced by the lack of a significant difference in the stop frames detected by the program. Samples sizes are as follows: All groups, n = 72. Significance was assessed using Kruskal Wallis test start frame groups, and Mann-Whitney U-test was used for end frame group. Significant differences are denoted as: *\**, *p* < 0.05.

### Analysis of tracking data

Another challenge faced by researchers utilizing the larval motor touch response assay is the analysis of the large amounts of tracking data (*e.g.,* drug or genetic screens). Conventionally, this required the manual import of output data into a spreadsheet for each video recorded. Then, researchers must manually add formulas to each video’s data in order to generate four swim metrics: swim duration (s), swim distance (mm), mean swim velocity (mm/s) and maximum swim velocity (mm/s) into complete datasets. While less challenging than tracking individual larval touch-responses, this post-tracking analysis requires time. As such, we included built-in analysis of tracking data as a feature of ZebraTrack. The program utilizes the user-specified value for x-y calibration (x-y cal) to relate pixel space to real space. The x-y cal value can be obtained by measuring an object in frame (*e.g.,* the diameter of the petri dish) using the measuring tool in ImageJ and dividing the length in millimetres of the object by its length in pixels. Thus, the user obtains a x-y cal value in units of millimetre-per-pixel. The ZebraTrack program functions under the assumption that all the videos in a batch are recorded with the same x-y cal value. In instances when the x-y cal changes (*e.g.,* distance of the camera lens to the petri dish) users will have to change the x-y cal value by starting analysis of a separate batch of videos. The program calculates swim duration by dividing the total number of frames by the frame rate. Swim distance is calculated by taking the sum of all cells in the “distance_since_last_frame” column, which is generated by taking the cartesian distance between the current and previous points and multiplying the result by the x-y cal value. Mean swim velocity is calculated by dividing the “distance_since_last_frame” value by the frame rate, for each cell, and then averaging all values in the swim pattern to obtain the mean swim velocity. Maximum swim velocity is calculated by finding the maximum value in the “velocity” column. In the final step, ZebraTrack takes each of the four swim metrics for each larva in the batch, and generates a summary page. This page shows each larva’s four swim metrics, as well as the mean for each metric across all larvae in the batch and outputs an .xlsx file.

### Detection of larval phenotypes

In order to determine the ability of the ZebraTrack program to detect subtle larval phenotypes, we conducted a simple assay in which zebrafish larvae were subjected to various temperatures allowing us to investigate the effect of temperature on motor function. We selected three temperatures to investigate. Larvae were raised at the normal rearing temperature of 28.5 °C but 1 hr prior to the motor touch assay larvae were switched to system water at a temperature of either 23.5 °C, 28.5 °C, or 33.5 °C. We hypothesized that elevated water temperature would result in larvae displaying increased swim duration (s), swim distance (mm), mean swim velocity (mm/s) and maximum swim velocity (mm/s). Whereas, decreased water temperature would result in lower values of our four motor function metrics. While swim duration was not significantly increased altered in the three temperature groups (**Fig. 4A**), we observed that swim distance was significantly increased in both the 28.5 °C and 33.5 °C groups when compared to the 23.5 °C group **(Fig. 4B)**. In addition, ZebraTrack detected an increase in mean swim velocity in larvae incubated at 33.5 °C when compared to larvae incubated at 23.5 °C **(Fig. 4C)**. Furthermore, ZebraTrack also detected an increase in maximum swim velocity in the 28.5 °C and 33.5 °C groups when compared to the 23.5 °C group **(Fig. 4D)**. Taken together, these results suggest that ZebraTrack is robustly capable of detecting subtle changes in larval motor touch-responses.

**Figure 4.**
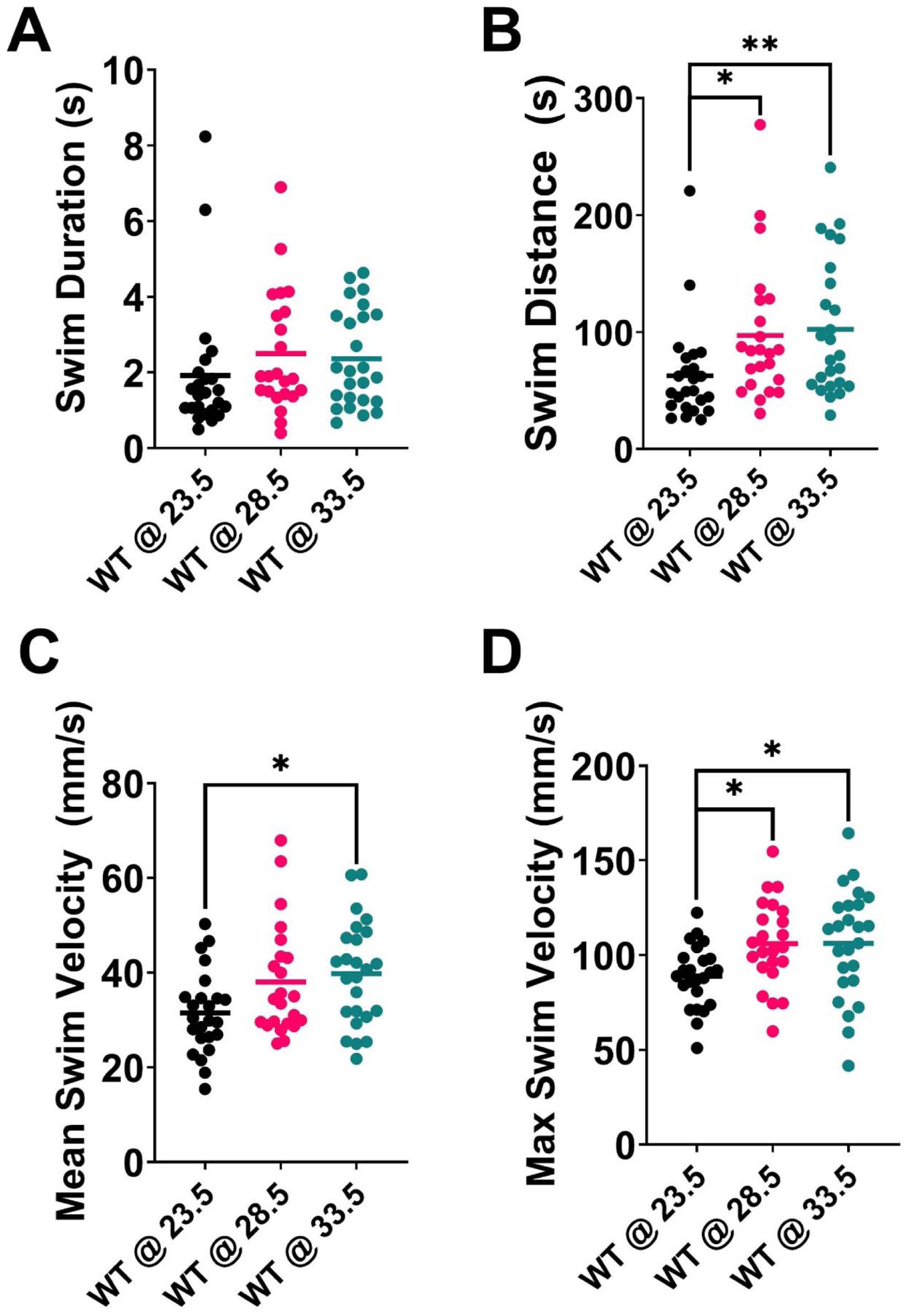
ZebraTrack captures temperature-dependent alterations in motor function of 2-day-old larval touch response videos. **A**, No statistically significant increases in swim duration were observed using ZebraTrack. **B**, Swim distance derived from ZebraTrack datasets captured that larvae incubated both the 28.5 °C (*p* = 0.018) and 33.5 °C (*p* = 0.01) were significantly increased when compared to larvae held at 23.5 °C. **C**, ZebraTrack identified a statistically significant increase in mean swim velocity in the 33.5 °C larval group when compared to larvae held at 23.5 °C (*p* = 0.027). **D**, Zebratrack identified a statistically significant increases in maximum swim velocity in larvae held at 28.5 °C (*p* = 0.035) and 33.5 °C (*p* = 0.03) when compared to larvae held at 23.5 °C. Samples sizes are as follows: 23.5 °C group, n = 24; 28.5 °C, n = 23; 33.5 °C, n = 25. Significance was assessed using Kruskal Wallis test for swim duration, swim distance and mean swim velocity, and One-Way ANOVA was used for maximum swim velocity. Significant differences are denoted as: *\**, *p* < 0.05; **, *p* < 0.01.

### Performance compared to manual tracking

To compare our findings obtained using ZebraTrack, we analyzed the temperature ramp touch-response data by hand using the ImageJ’s manual tracking plugin. While manual analysis is subject to experimenter bias and error, it remains a useful tool for comparison. We hypothesized that ZebraTrack’s results would capture similar findings in our datasets. Manual tracking, similarly to ZebraTrack, did not reveal any significant differences in swim duration across the three temperature groups **(Fig. 5A)**. Manual tracking also shows statistically significant increases in swim distance in the 28.5 °C and 33.5 °C groups when compared to the 23.5 °C group **(Fig 5B)**. Notably, manual tracking did not reveal any significant differences in mean swim velocity **(Fig. 5C)**. We observed a statistically significant increase in maximum swim velocity in both the 28.5 °C and the 33.5 °C groups when compared to the 23.5 °C group **(Fig. 5D)**. Taken together, these results demonstrate that all significant differences derived from using the ImageJ manual tracking plugin were also observed using ZebraTrack. Furthermore, to ascertain the differences between manual analysis and machine learning-based tracking, we compared manual tracking data to data obtained using ZebraTrack for the 28.5 °C group (**Fig. 6**). We observed no statistically significant differences between the two datasets, indicating that ZebraTrack adequately captures larval motor touch responses metrics. While both swim duration (**Fig. 6A**) and swim distance (**Fig. 6B**) had similar values from manual analysis to machine learning analysis, we observed a slight trend of decreased mean swim velocity values from manual analysis when compared to the data captured by ZebraTrack (**Fig. 6C**). Conversely, we also observed a slight trend of increased maximum swim velocities from manual tracking when compared to the data captured by ZebraTrack (**Fig. 6D**). In summary, these findings indicate that the data captured by ZebraTrack does not deviate significantly from manual tracking data, indicating its robustness as a replacement for labor-intensive ImageJ’s manual tracking.

**Figure 5.**
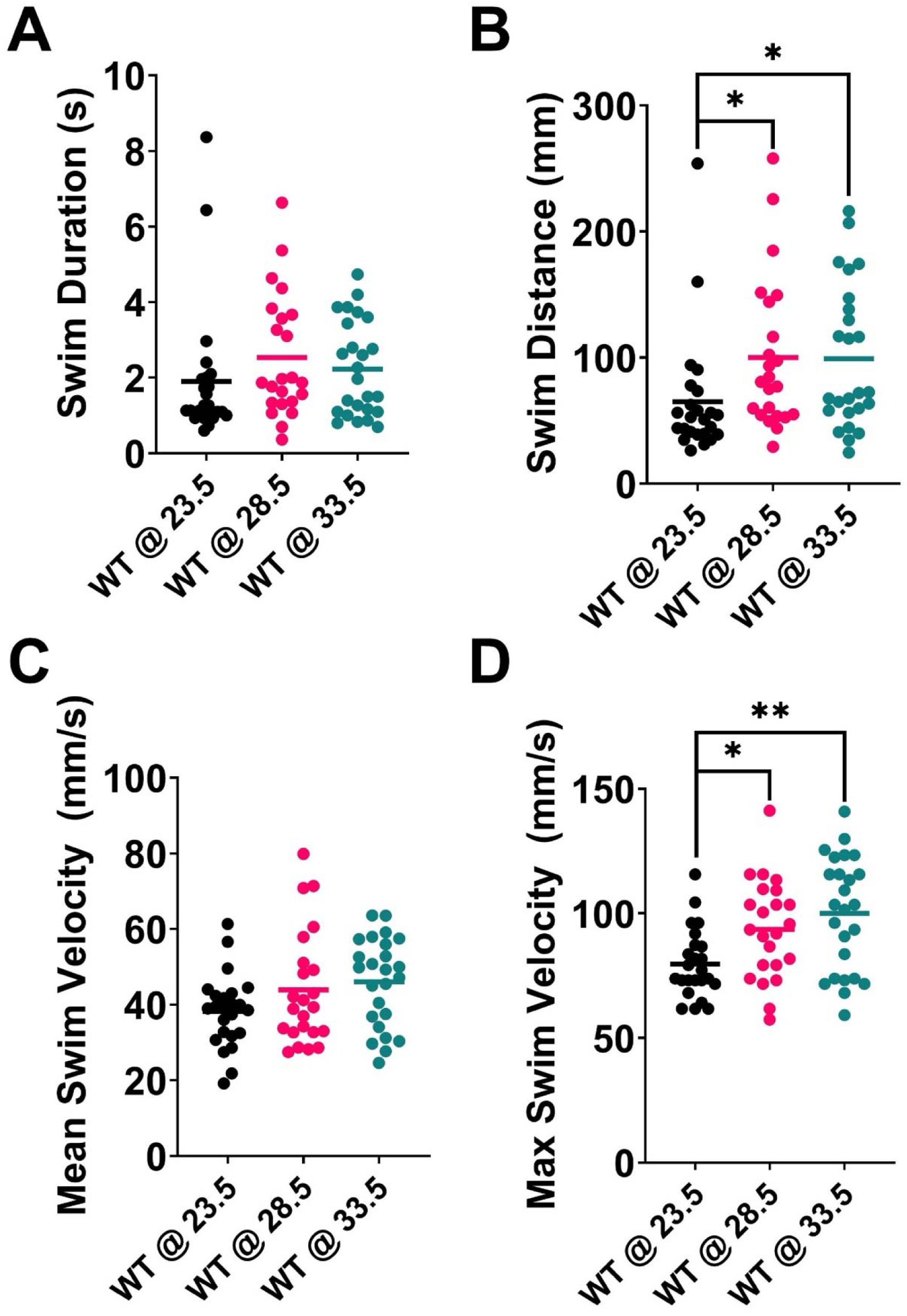
Manual analysis using the ImageJ Manual Tracking plugin captures temperature-dependent alteration of larval motor function from the same data as in Figure 4. **A**, No significant differences in swim duration were observed across the three temperature groups. **B**, Larvae incubated at both the 28.5 °C (*p* = 0.019) and 33.5 °C (*p* = 0.023) displayed significantly increased swim distance when compared to larvae held at 23.5 °C. **C**, Manual analysis did not reveal any statistically significant differences in mean swim velocity. **D**, Statistically significant increases in maximum swim velocity in larvae held at 28.5°C (*p* = 0.042) and 33.5 °C (*p* = 0.001) were observed when compared to larvae held at 23.5 °C. Samples sizes are as follows: 23.5 °C group, n = 24; 28.5 °C, n = 23; 33.5 °C, n = 25. Significance was assessed using Kruskal Wallis test with post-hoc Dunn’s test for swim duration, swim distance and mean swim velocity, and One-Way ANOVA with post-hoc Tukey’s test was used for maximum swim velocity. Significant differences are denoted as: *\**, *p* < 0.05; **, *p* < 0.01.

**Figure 6.**
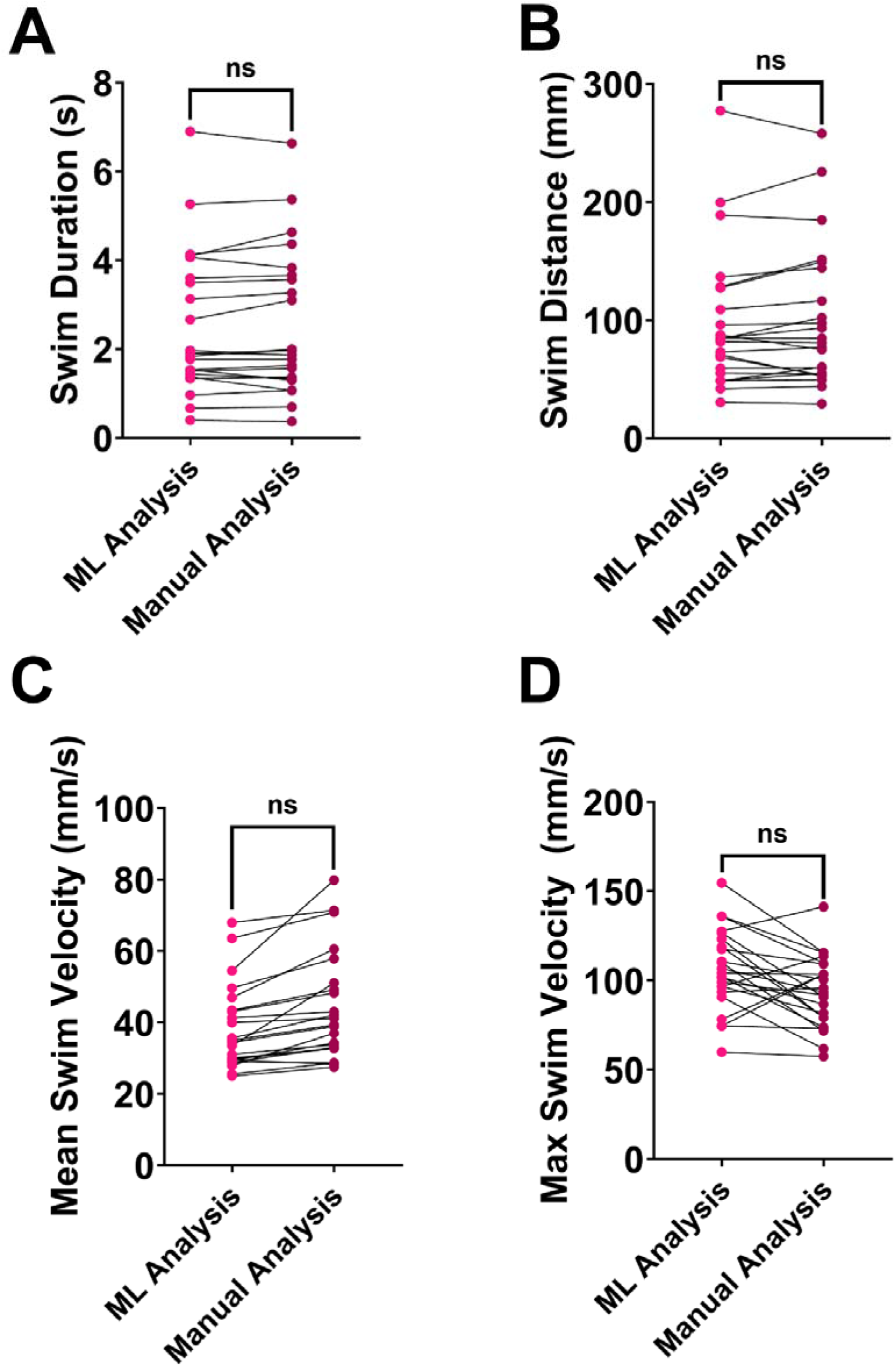
Machine learning analysis does not significantly alter data compared to manual analysis. The data presented in this figure is from the 28.5 °C group, as analyzed manually and by the ZebraTrack program. **A**, Machine learning analysis does not significantly alter the data obtained for swim duration when compared to manual analysis. **B**, Machine learning analysis does not significantly alter the data obtained for swim distance when compared to manual analysis. **C,** Machine learning analysis tends to decrease the values for mean swim velocity when compared to manual analysis, but does not significantly alter the data. **D**, Machine learning analysis tends to increase the values for maximum swim velocity when compared to manual analysis but does not significantly alter the data. Samples sizes are as follows: 28.5 °C group, n = 23. Significance was assessed using Mann-Whitney U-test for swim duration, swim distance and mean swim velocity, and Welch’s t-test for maximum swim velocity. No significant differences were identified.

## Discussion

In this study, we introduce ZebraTrack, an accessible, fully automated pipeline for the analysis of two-day old larval zebrafish motor touch-response videos. Using a custom-trained YOLOv8n object detection model, combined with bespoke error correction and post-tracking analysis, the program allows for accurate high-throughput analysis of motor touch-response data without the need for expensive software or extensive manual labor. Our results demonstrate that ZebraTrack performs comparably to manual tracking larval swim metrics across experimental groups, validating its use as a reliable tool for behavioural phenotyping.

A key strength of ZebraTrack lies in its scalability and accessibility. The use of Google’s Colaboratory platform and free file-hosting *via* Google Drive ensures that researchers without access to expensive specialized hardware or proprietary software can still efficiently perform large-scale behavioral analyses. Moreover, the use of the YOLOv8n model further reinforces this accessibility, as it allows for rapid analysis of motor touch response videos even on modest computing workstations. While larger models may offer marginal gains in accuracy, our results suggest that YOLOv8n provides sufficient precision for the relatively simple task of single object tracking in standardized video conditions.

ZebraTrack serves as an example that even with limited data, it is possible to achieve high performance on a computer vision task using the YOLOv8n model. While we hypothesized that performance (as measured by mAP50-95 on separate, unseen test datasets) would increase with larger amounts of training data, we observed the opposite trend; the mAP50-95 value for the first dataset of 18,189 images was 0.74. With an increase to 20,014 images for the second dataset, a mAP50-95 value of 0.79 was achieved. However, on the final dataset of 65,416 images, we observed a decrease in mAP50-95 to a value of 0.74. There are multiple interpretations of this decrease in performance. For example, it is possible that the largest dataset, which contained a wider range of data, was more variable than smaller datasets. Thus, it may have captured a more realistic picture of the variability inherent to larval touch response videos collected by different researchers. For example, various distractor objects such as hair, hands, and forceps may occasionally occlude larva in videos, larval pigmentation may vary based on their genotype or treatment, and experimenters may have slightly different techniques for evoking the motor responses, resulting in different situations for the model to interpret. By providing a larger dataset, we provide a more realistic snapshot of what future videos may look like, thus providing the model with a more robust training scope despite the slight reduction in overall performance. Conversely, it could be argued that a smaller amount of training data was sufficient to achieve usable performance for this task, and that adding more data was not beneficial to performance. Nevertheless, our findings suggest that the pre-training provided with YOLOv8n is sufficient to offset limited datasets, allowing this model to be used even in circumstances when researchers would like to implement a machine learning solution without an abundance of training data.

To demonstrate the utility of ZebraTrack, we examined the effect of temperature on motor touch-responses and compared these to manual tracking results obtained from the same datasets. We observed that ZebraTrack reliably detects the subtle differences in motor function metrics in larval touch responses, as evidenced by its ability to detect similar trends to the current standard of tracking larval motor patterns (ImageJ’s manual tracking). While the two methods for tracking metrics of motor function were similar, we observed a statistically significant increase in mean swim velocity from the 23.5 °C group to the 33.5 °C group in ZebraTrack’s results, where manual analysis failed to reveal any such difference (**Figure 4C** & **Figure 5C**). Given these results, we theorized that the ability of ZebraTrack to robustly track a single location on the larvae (*i.e.*, the larval head) provides more accurate, less noisy tracking data compared to error-prone manual analysis, thus reducing Type II errors.

During the development of ZebraTrack, two issues routinely encumbered the program, which were the detection of larval swim pattern initiation and termination (henceforth referred to as larval starts and larval stops, respectively). To overcome these issues, we integrated a One Euro filter and a velocity thresholding strategy, thereby allowing ZebraTrack to successfully identify the start and end of larval zebrafish swim patterns (**Figure 3**). The One Euro filter is commonly used in applications requiring real-time tracking, such as motion capture, human-computer interaction, and robotics (Casiez et al., 2012). While our application does not involve real-time tracking, the same principles apply when capturing larval swim patterns. The thresholding approach described herein robustly detects larval starts and stops but is subject to certain limitations. On one hand, the detection of larval starts can be mired by the presence of distractor objects lowering tracking fidelity and thus disrupting our thresholding approach. Whereas on the other hand, detection of larval stops is complicated by their inherent variability, owing to the gradual larval deceleration and the noisier nature of tracking data at low speeds. While these limitations mirror the ambiguity often encountered in manual analysis, improving the larval start and stop detection algorithm represents a direction for future iterations of algorithmic approaches to tracking larval swim patterns. Incorporating machine learning-based temporal classifiers or recurrent models could help increase the reliability of start and stop detections compared to the thresholding approach currently taken by ZebraTrack.

Importantly, ZebraTrack’s ability to distinguish temperature-dependent differences in swim metrics demonstrates its practical utility in behavioural assays. The detection of statistically significant changes in swim distance, mean velocity, and maximum velocity in larvae exposed to elevated temperatures is consistent with known effects of temperature on zebrafish physiology (Herbing, 2002; Green and Fisher, 2004). These findings were further corroborated by manual tracking, which revealed similar patterns to the tracking data provided by ZebraTrack. Our results support the use of ZebraTrack as a robust, user-friendly, and cost-effective alternative to the ImageJ’s manual tracking. Given that the start/stop detection capabilities of the program remain the main limitation of ZebraTrack, albeit a minor limitation, we suggest that users of ZebraTrack manually review the data output by the program. Nonetheless, the benefits of ZebraTrack’s automation, consistency, and accessibility make it a valuable tool for the zebrafish research community. Future developments could include integration with graphical user interface, improved larval start and stop detection, or real-time behavioral classification. Ultimately, ZebraTrack dramatically reduces analysis time for the motor touch response assay from hours to seconds, and thus it may accelerate the pace of discovery in future studies using zebrafish.

## Data availability statement

The code and data underlying this research are available on GitHub at [https://github.com/Armstrong-Lab-70/ZebraTrack] as well as on Figshare at [https://dx.doi.org/10.6084/m9.figshare.29631308]. Please note that, due to file size restrictions, our training datasets for YOLOv8n cannot be published online but will be shared upon reasonable request.

## Acknowledgements

The authors would like to thank Dr. Abel González Garcia for helpful discussions and mentorship in computer vision techniques.

## Supplemental Video 1

**Supplemental Video 1.** An example video of a larval zebrafish exhibiting the touch-evoked motor response, overlayed with YOLOv8n’s tracking predictions.

## References

Albadri S, De Santis F, Di Donato V, Del Bene F (2017) CRISPR/Cas9-Mediated Knockin and Knockout in Zebrafish. In: Genome Editing in Neurosciences (Jaenisch R, Zhang F, Gage F, eds), pp 41–49. Cham (CH): Springer

Armstrong GA, Drapeau P (2013) Calcium channel agonists protect against neuromuscular dysfunction in a genetic model of TDP-43 mutation in ALS. J Neurosci 33:1741–1752.

Brustein E, Saint-Amant L, Buss RR, Chong M, McDearmid JR, Drapeau P (2003) Steps during the development of the zebrafish locomotor network. Journal of Physiology-Paris 97:77–86.

Casiez G, Roussel N, Vogel D (2012) 1 € filter: a simple speed-based low-pass filter for noisy input in interactive systems. In: Proceedings of the SIGCHI Conference on Human Factors in Computing Systems, pp 2527–2530. Austin, Texas, USA: Association for Computing Machinery.

Chumachenko K, Gabbouj M, Iosifidis A (2022) Chapter 11 - Object detection and tracking. In: Deep Learning for Robot Perception and Cognition (Iosifidis A, Tefas A, eds), pp 243–278: Academic Press.

Goldshtein H, Muhire A, Petel Légaré V, Pushett A, Rotkopf R, Shefner JM, Peterson RT, Armstrong GAB, Russek-Blum N (2020) Efficacy of Ciprofloxacin/Celecoxib combination in zebrafish models of amyotrophic lateral sclerosis. Annals of Clinical and Translational Neurology 7:1883–1897.

Green BS, Fisher R (2004) Temperature influences swimming speed, growth and larval duration in coral reef fish larvae. Journal of experimental marine biology and ecology 299:115–132.

Herbing IHv (2002) Effects of temperature on larval fish swimming performance: the importance of physics to physiology. Journal of Fish Biology 61:865–876.

Kimmel CB, Ballard WW, Kimmel SR, Ullmann B, Schilling TF (1995) Stages of embryonic development of the zebrafish. Developmental Dynamics 203:253–310.

Masino MA, Fetcho JR (2005) Fictive Swimming Motor Patterns in Wild Type and Mutant Larval Zebrafish. Journal of Neurophysiology 93:3177–3188.

Patten SA, Aggad D, Martinez J, Tremblay E, Petrillo J, Armstrong GA, La Fontaine A, Maios C, Liao M, Ciura S, Wen XY, Rafuse V, Ichida J, Zinman L, Julien JP, Kabashi E, Robitaille R, Korngut L, Parker JA, Drapeau P (2017) Neuroleptics as therapeutic compounds stabilizing neuromuscular transmission in amyotrophic lateral sclerosis. JCI Insight 2.

Saint-Amant L, Drapeau P (1998) Time course of the development of motor behaviors in the zebrafish embryo. Journal of Neurobiology 37:622–632.

Saint-Amant L, Drapeau P (2000) Motoneuron Activity Patterns Related to the Earliest Behavior of the Zebrafish Embryo. The Journal of Neuroscience 20:3964–3972.

Simone BW, Martínez-Gálvez G, WareJoncas Z, Ekker SC (2018) Fishing for understanding: Unlocking the zebrafish gene editor’s toolbox. Methods 150:3–10.

Sohan M, Ram T, Ch V (2024) A Review on YOLOv8 and Its Advancements. In, pp 529–545.

Stewart AM, Braubach O, Spitsbergen J, Gerlai R, Kalueff AV (2014) Zebrafish models for translational neuroscience research: from tank to bedside. Trends in Neurosciences 37:264–278.

Sztal TE, Ruparelia AA, Williams C, Bryson-Richardson RJ (2016) Using Touch-evoked Response and Locomotion Assays to Assess Muscle Performance and Function in Zebrafish. J Vis Exp.

Talha MM, Khan HU, Iqbal S, Alghobiri M, Iqbal T, Fayyaz M (2023) Deep learning in news recommender systems: A comprehensive survey, challenges and future trends. Neurocomputing 562:126881.

Varghese R, Sambath M (2024) YOLOv8: A Novel Object Detection Algorithm with Enhanced Performance and Robustness. In: 2024 International Conference on Advances in Data Engineering and Intelligent Computing Systems (ADICS), pp 1–6.

Varshney GK, Sood R, Burgess SM (2015) Understanding and Editing the Zebrafish Genome. Adv Genet 92:1–52.

Westerfield M (2000) The zebrafish book : a guide for the laboratory use of zebrafish (Danio rerio), Ed. 5. Edition. Eugene, Or.: M. Westerfield.

Yaseen M (2024) What is YOLOv8: An In-Depth Exploration of the Internal Features of the Next-Generation Object Detector.

Zaidi SSA, Ansari MS, Aslam A, Kanwal N, Asghar M, Lee B (2022) A survey of modern deep learning based object detection models. Digital Signal Processing 126:103514.

